# Cell division dynamics generate heterogeneous contact-mediated signaling outputs

**DOI:** 10.64898/2026.06.18.733180

**Authors:** Jonathan E. Dawson, Abdul Naseer Malmi-Kakkada

**Affiliations:** Engineering and Physics Department, Whitworth University, Spokane, WA 99251, USA; Department of Physics, Astronomy and Geosciences, Towson University, Baltimore, MD 21252, USA

## Abstract

Contact mediated cell-cell communication where direct physical contact between adjacent ligand cells and receptor cells trigger signal output is important during growth, development and regeneration of organisms. While the molecular machinery underlying contact mediated cell signaling is well explored, how the local spatial context of cells affect cell-cell contact mediated gene expression is not clear. Here, we present a vertex-based computational model to study spatial and temporal behavior of contact mediated signal output (which we refer to as output) in growing cell collectives. We consider cell-cell contact length dependent output synthesis and output degradation in receptor cells together with cell division to understand how dynamics at the scale of single cells lead to heterogeneous signal output. By tracking single receptor cells over time in growing cell collectives *in silico*, we show that cell growth and division lead to continuous and dynamic rearrangement of cell-cell contact between receptor and ligand cells which in turn affect the output levels. Our model predicts that the orientation of cell division plays a key role in the heterogeneity of signal output. We elucidate the link between cell mechanical properties that control cell shape, growth, and division, with signal output in receptor cells during contact mediated signaling processes.

## I. INTRODUCTION

Cell-cell interactions (CCIs) are necessary for multicellular life to exist, allowing cells to execute collective functions and coordinate their activities [1–5]. Generally, cells interact with one another to activate signalling pathways in other cells [2], coordinate gene expression [6] and drive cell functions [7]. While these interactions may involve a multitude of components such as proteins, small compounds and the extracellular matrix etc, ligands that bind with cognate receptors on other cells is the dominant mechanism used for CCIs [8]. The importance of CCIs in governing tissue physiology [9], disease [10] and development [11] is now becoming apparent.

Computational tools have helped unravel important CCI modalities and new tools are emerging to address complex properties of intercellular interactions [3, 8]. Experimentally, major approaches in studying CCIs include single-cell transcriptomics [12], droplet based methods [13], cell-cell contact dependent chemical labeling [14] and synthetic circuits [15, 16]. Here, our focus is on a synthetic form of juxtacrine signaling that operates orthogonally to native processes and can be precisely controlled, making them a powerful reductionist tool with which to address fundamental questions in CCIs [17]. Previously, we investigated how cell-cell contact length and tissue growth dynamics affect juxtacrine signal responses by implementing a custom synthetic gene circuit (synthetic Notch or synNotch) in *Drosophila* wing imaginal discs alongside mathematical modeling [16]. We discovered that the area of contact between cells largely determines the extent of synNotch activation with the output forming a graded spatial profile that extends several cell diameters from the signal source [16]. This provided evidence that the response to juxtacrine signals can persist in cells as they proliferate away from source cells. Our experimentally verified computational model predicted spatially graded outputs without diffusion or long-range cell-cell communication [16].

By building on our previous work, here we focus on how time-dependent juxtacrine signal output is affected by cell-cell contact parameters in a growing tissue. Given the importance of cell division in the generation of cell diversity [18–21] and the importance of juxtacrine signaling during development [22, 23], we sought to understand how cell division patterns influence juxtacrine signal output and lead to heterogeneous output patterns.

Contact-mediated signaling, such as the ubiquitous Notch signaling, occurs through physical contact between a ligand (‘signal sending’) cell and a receptor (‘signal receiving’) cell [24, 25]. A series of steps then lead to target gene activation (which we refer to as ‘output’) in the receptor cell, Fig. 1(A). Although contact-mediated signaling regulates crucial cell-fate decisions at an individual cell level and coordinates spatio-temporal patterning in tissues [26], little is known about how cell division dynamics impact output dynamics. During tissue growth, cells grow in size, divide and undergo spatial rearrangement within the tissue. Hence, the cell-cell contact length responsible for contact-mediated signaling [16, 27] change continuously over time, Fig. 1(B)-(C). We showed previously [16] that this process leads to two types of receptor cells: (i) peripheral cells that maintain contact with ligand cells (see cells with borders marked in red in Fig. 1(C)) and (ii) interior cells that do not have direct contact with ligand cells. Since the signal output is generated actively only in the peripheral receptor cells, where they are in contact with the ligand cells, it remains to be understood how variations in the individual cell-cell contact lengths due to cell growth and division affect the signal output; Fig. 1(C) (see bottom panel).

**Figure 1.**
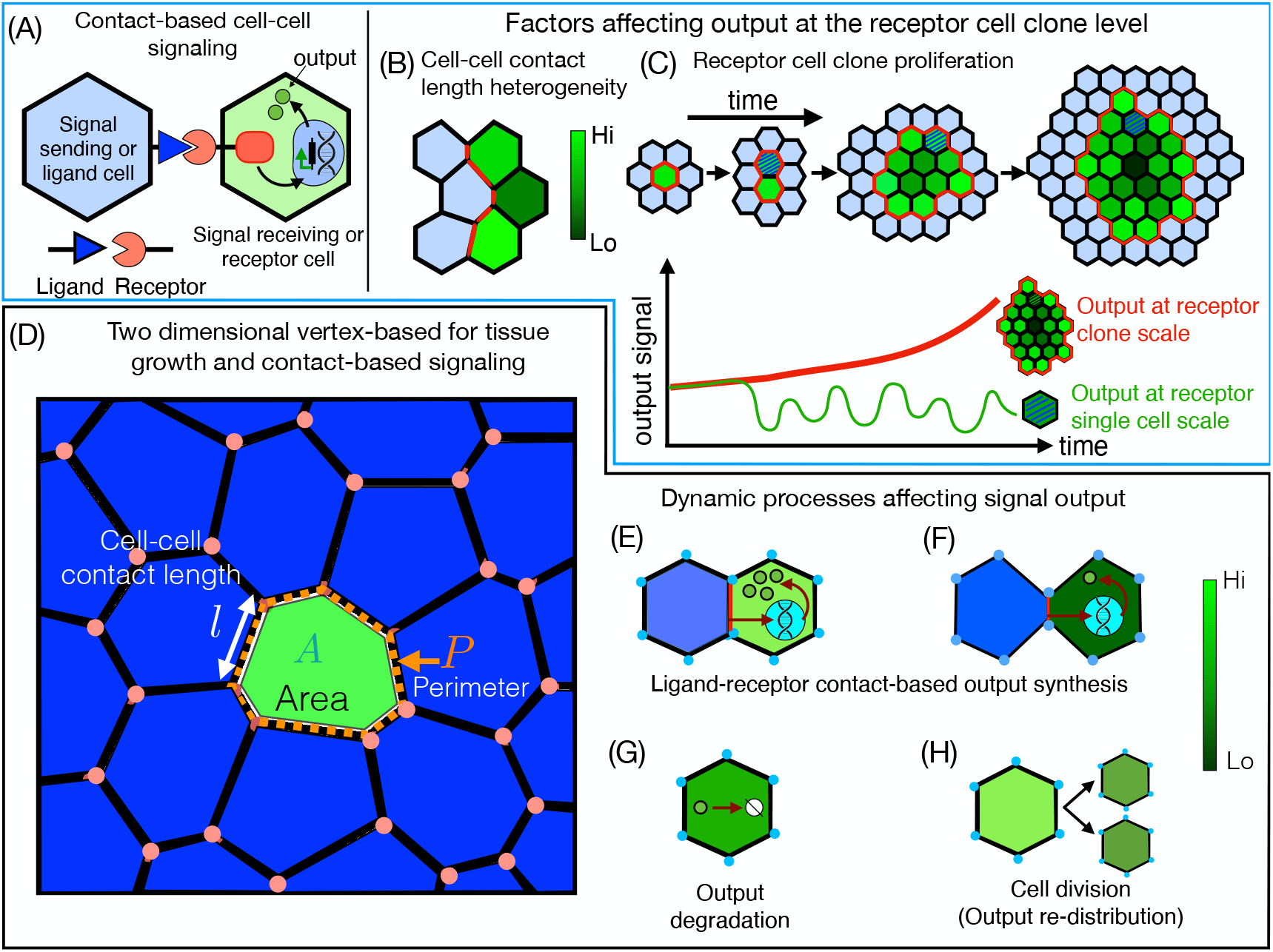
Dynamics of contact-based intercellular signaling. (A) When a membrane-anchored ligand of a signal-sending cell, also known as a ligand cell, engages with a receptor on the surface of an adjacent signal-receiving cell, also known as a receptor cell, a signal, in the form of an output, is elicited in the receptor cell. We consider a synNotch circuit with green fluorescent protein (GFP) output which allows for visualization and quantification of the output. (B) The strength of the output in a receptor cell depends on several factors, one of those being the contact length shared between the receptor cell and the ligand cell. (C) In a growing cluster of signal-receiving cells, also known as the receptor clone, the cell-cell contact length continuously changes due to cell division, which affects the output strength and its spatial distribution within the receptor clone. As a result, although the strength of individual cell output within the clone might fluctuate highly over time, the total output of the receptor clone could show a monotonic behavior (bottom panel). (D-F) A two-dimensional vertex model is used to study the dynamics of contact-based signaling in growing epithelial tissue. The model integrates active cell growth and division with intracellular output biosynthesis, which depends on the cell-cell contact length shared between the receptor cell (green colored) and the ligand cells (blue colored). (G) The model accounts for output degradation in the receptor cells over time. (H) When a receptor cell divides in the model, the output is equally distributed between the two resulting daughter cells.

Our work and those of others [16, 27–29] have pointed to the coordination between juxtacrine signal output and the biophysical properties of the cell-cell contact interface in obtaining heterogeneous signal output levels. Here, we show how cell division orientation with respect to the contact plane between signal sending and signal receiving cells can give rise to marked differences in the output in the newly divided cells (or daughter cells). Even with consistent cell size changes across cell division cycles, cell-cell contact length markedly varies in between and over consecutive cell division cycles, leading to highly heterogenous output levels.

## II. MODEL DESCRIPTION AND SIMULATION DETAILS

To understand the effect of cell growth and division on juxtacrine signal output patterns, we developed a vertex-based mathematical model in proliferating tissues, Fig. 1(D). In our model, cells are represented as a network of interconnected two-dimensional polygons [30, 31]. Each cell in this framework is characterized by an energy function that describes the balance between cell volume incompressibility, active contractility due to actomyosin cortex, and cell-cell adhesion [30] (see Appendix for more details). We introduce a chemomechanical coupling between intracellular signal output (simply referred to as output hereafter) dynamics and vertex-based cell shape dynamics, including cell growth, and division. The *in silico* tissue (Fig. 1(D)) consists of two cell types: (i) the receptor cells that produce the output (Fig. 1(D), green) and (ii) the ligand cells (Fig. 1(D), blue) that signal to adjacent receptor cells. The output strength in receptor cells is represented by a green color scale, with black color representing no signal output and bright green color representing high signal output (Fig. 1(E, F)). We hypothesize that the rate of change of signal output (*G*_*α*_ [arbitrary units]) with time (*t*) is governed by the dynamic equation [16],

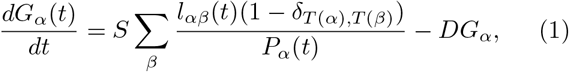

where, *l*_*αβ*_ is the time-dependent cell-cell junction contact length, *P*_*α*_ is the perimeter of the receptor cell *α* = 1, 2, 3…*N*, with *N* being the total number of receptor cells. The summation is over all the cells *β* in contact with a cell *α*. The signal output is synthesized only in those receptor cells that are in contact with the ligand cells, as enforced by the delta function term (see Apendix for further details). The first term on the righthand side of Eq. (1) describes the output synthesis at the rate *S* [*h*^−1^] (Fig. 1(E), *h*^−1^ denotes 1*/*hour), and increases with larger contact lengths between receptor and ligand cells (Fig. 1(E, F)) [32]. The second term on the right hand of Eq.(1) accounts for the output degradation in the receptor cell *α* at a constant rate of *D* (h^−1^) (Fig. 1(G)). A key feature of our model is that the output is proportional to the instantaneous cell-cell contact length shared directly between the receptor and ligand cells. As the cell-cell junction contact length can vary as a function of time due to growth-driven cell shape changes and cell division [36], our model captures the effect of intrinsic cellular level processes on output during tissue growth.

As cell growth and division are important parameters in our study, we implement specific rules for such cell behaviors. A cell divides along its short axis when its area doubles, resulting in tissue growth over time (see Appendix for details). Upon division, the total amount of signal output in a parent cell is distributed equally between the two resulting daughter cells (Fig. 1(H)). The simulation initial conditions and the relevant parameter values are listed in Table I and described further below.

**Table 1.** Model parameters, their meaning, dimensionless values and equivalent magnitude in experimental units.

| Model parameter | Description | Values (Dimensionless) | Experimental values |
| --- | --- | --- | --- |
| $A_b$ | Cell base area | 50 | $\sim 10 \mu m^2$ [33] |
| $A_0$ | Range of cell preferred area | [50-100] | $[10 - 20] \mu m^2$ |
| $P_0$ | Cell preferred perimeter | 25 | $\sim 11 \mu m$ [33] |
| $K_A$ | Area elastic constant | 500 | |
| $K_P$ | Perimeter elastic constant | 100 | |
| $g_0$ | Cell area growth rate | 5 | $1 \mu m^2/h$ |
| $S$ | Rate of cellular GFP output | 2 | $0.5 h^{-1}$ |
| $D$ | Rate of cellular GFP degradation | [0-0.5] | $\sim [0-0.5] h^{-1}$ [34, 35] |
| $dt$ | Time step | 1 | 1 h |

### Initial conditions

We initiated the simulations with 16 ligand cells and 1 receptor cell. Initially, the receptor cell has no signal output (*G* = 0 at *t* = 0 h). With time the dynamics of the receptor cell output is determined by Eq. 1. Cell preferred area was fixed at ~ 10*µm*^2^ with each cell growing at the same rate *g*_0_ = 1 *µm*^2^*/h*. We keep all the parameters fixed, except for the output degradation rate *D* (see Table I). We recorded, at each time point, the cell-cell contact length, perimeter and the output (*G*_*α*_) of each receptor cell situated at the interface between ligand and receptor clones. Each simulation was run for total time of 80 h, sufficient to account for multiple cell division cycles.

## III. OUTPUT IN THE RECEPTOR CELLS EXHIBIT HETEROGENEITY IRRESPECTIVE OF SYNTHESIS AND DECAY RATES

Most studies of juxtacrine signaling pathways consider signaling events at the scale of up to 2 cells in isolation [27] or at the level of an average cell that is assumed to represent the whole cell population [37]. Similarly, experimental output measurements are usually averaged over populations consisting of many hundreds or thousands of cells. This approach does not account for the fact that even within a homogeneous population of signal sending and receiving cells, local conditions may influence the output and result in phenotypic heterogeneity.

To investigate this further, we quantified the spatial and time-dependent behavior of the total output in the receptor clone. We begin with one receptor cell surrounded by ligand cells and let the tissue evolve to *t* = 80 h at which point there are ~ 70 cells in the receptor clone, Figs. 2(A)-(B). The dynamic equation that describes time-dependent changes in output within individual receptor cells is given by Eq.1. We observe a heterogeneous spatial distribution of the output in the receptor clone irrespective of the degradation rate, *D* = 0 *h*^−1^ and *D* = 0.5 *h*^−1^. Output is higher in the receptor cells at the outermost layer of the clone as compared to those in the interior of the clone, Figs. 2(A) and (B).

**Figure 2.**
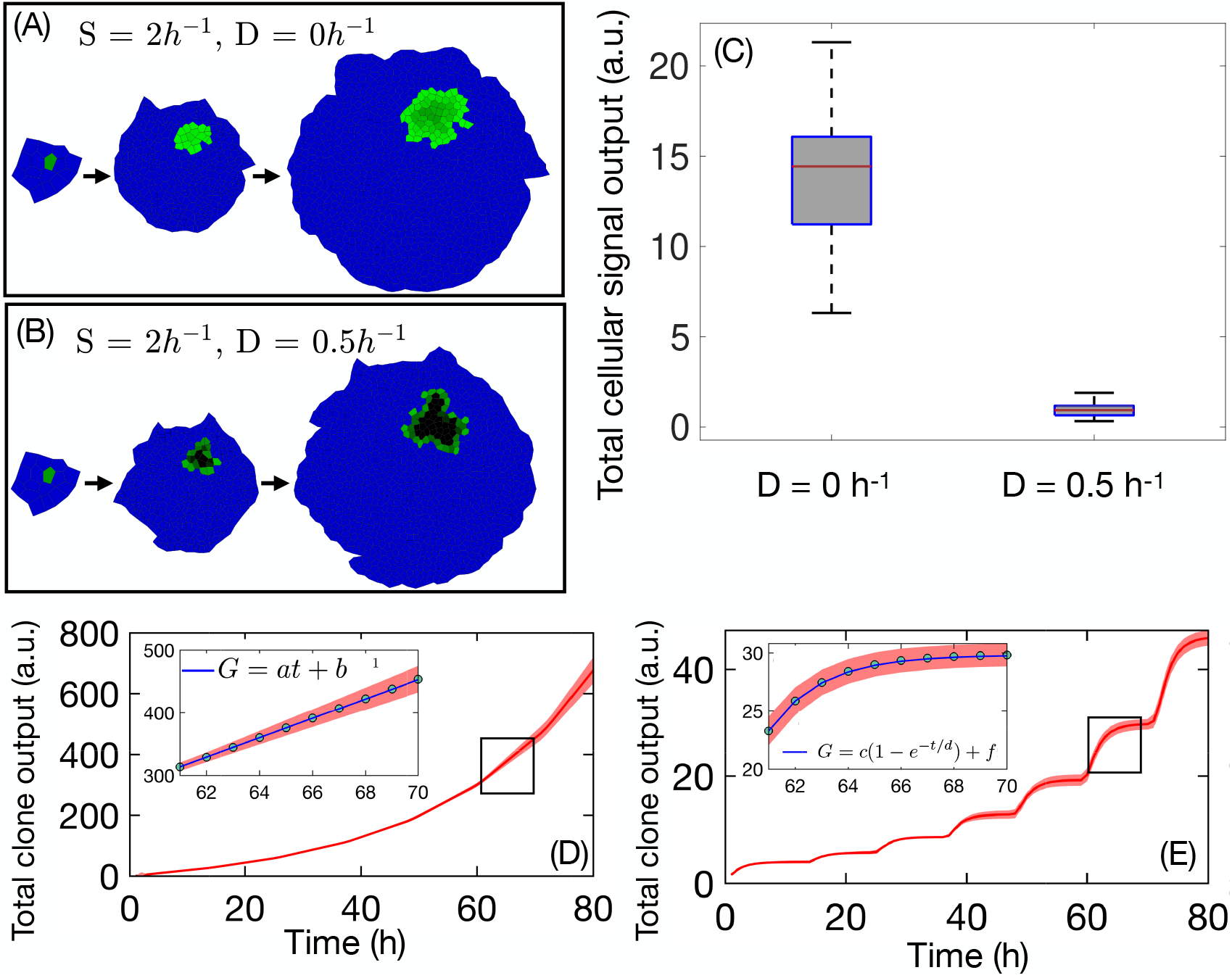
Spatial and time-dependent behavior of contact-based signaling output in a proliferating epithelial tissue. (A)-(B)Snapshots of a field of proliferating cells with one initial synNotch cell (green) surrounded by ligand cells (blue) at the initial time (*t*_0_) and at two other time points. Output synthesis (*S*) and degradation (*D*) rates are fixed to (A) *S* = 2 *h*^−1^, *D* = 0 *h*^−1^, and (B) *S* = 2 *h*^−1^, *D* = 0.5 *h*^−1^. (C) Even though the output is synthesized at a constant rate in all border receptor cells, there is considerable variation in the output without (*D* = 0 *h*^−1^) and with output degradation (*D* = 0.5 *h*^−1^). Mean values are denoted as a red line. All receptor cells situated at the border of the clone are in direct contact with the ligand cells, resulting in output synthesis at a constant rate *S* in all those receptor cells. (D) The output grows linearly with time for *D* = 0 *h*^−1^, as shown by a linear fit (inset) between two cell division events. (E) For *D* = 0.5 *h*^−1^, the total output tends to saturates between any two rounds of cell division, as shown by the fit to an exponential function (inset).

### Single cell output

To quantify the heterogeneity in the output levels, we focused on the receptor cells that are situated at the border of the clone and maintain active contact with ligand cells. By fixing the value of *S* and *D*, we ensured that the synthesis and degradation rates are uniform between all the cells within the tissue. Therefore, the variation in the output levels must then arise due to the difference in contact length and perimeter of the cells. When we measured the output levels in these border cells, we noticed considerable variation between cells (see Fig. 2(C)). For *D* = 0 *h*^−1^, the mean single cell output is = 13.9 (a.u.) with a standard deviation of = *±*3.9, while for *D* = 0.5 *h*^−1^, mean = 0.9 (a.u.) and standard deviation = *±*0.4. The coefficient of variation (CV) was calculated for each case (*D* = 0 *h*^−1^ and *D* = 0.5 *h*^−1^) as a relative measure of dispersion in the output values. We obtained values of CV in the range of 0.3 − 0.4 for both values of *D*.

### Total receptor clone output

Next, we studied the temporal behavior of the total output of the entire receptor clone, which is the sum of the individual receptor cell outputs. The total output receptor clones for both *D* = 0 *h*^−1^ and *D* = 0.5 *h*^−1^ shows an overall increase over time, Fig. 2(D)-(E). However, there is distinct difference in the temporal behavior of the output between successive rounds of cell division. For *D* = 0 *h*^−1^, focusing on the time window between two successive rounds of cell divisions, the average total output grows linearly with time (see Fig. 2(D-inset). The linear behavior of the total output was confirmed by fitting a straight line. On the other hand, when *D* = 0.5 *h*^−1^, there is a gradual saturation to a constant value which is well fit by an exponential function, Fig. 2(E-inset). To better understand these differing trends, we looked at the mathematical model given by Eq. 1. By summing Eq. 1 over each cell *α* in the receptor clone, we obtain the equation for the total output,

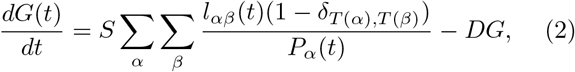

where, *G* = ∑_*α*_ *G*_*α*_. The first term in Eq. 2 contributes to the growth in the total output, whereas the second term accounts for the output degradation in the receptor clone. Our simulation results show that in the case of no output degradation, i.e., *D* = 0 *h*^−1^, total output grows linearly over time. According to our model, a linear time-dependent behavior of the clonal output, in the case of *D* = 0 *h*^−1^, is possible only if both the cell-cell contact length *l*_*αβ*_ and the cell perimeter *P*_*α*_ have the same functional dependence on time. In other words, *l*_*αβ*_(*t*)*/P*_*α*_(*t*) = *r*, where is *r* is an arbitrary constant, which gives the analytical solution of *G* = *Srt*, Fig. 2(D). Assuming *l*_*αβ*_(*t*)*/P*_*α*_(*t*) is a constant, as suggested by the linear behavior of *G* in the case of *D* = 0 *h*^−1^, Eq. 2 can also be analytically solved for *D* = 0.5 *h*^−1^ to obtain 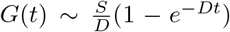, which is exactly the form of the function used to fit the simulation result of clonal output as a function of time, Fig. 2(E).

## IV. TIME-DEPENDENT BEHAVIOR OF CELL-CELL CONTACT LENGTH AT THE CLONAL INTERFACE IS HIGHLY HETEROGENEOUS

Because the output level in the receptor cell depends on the length of contact with ligand cells [16, 27], we wondered how time-dependent changes in cell-cell contact length affects the output. We anticipate the cell-cell contact length to increase with time between cell division events due to cell growth. However, the contact length may exhibit more abrupt time-dependent changes due to cell division. In epithelia, cells typically round up, constrict in the middle and divide symmetrically leading to significant changes in shape and contact length with neighboring cells [38]. Here, we focus on the scenario where a cluster of receptor cells together form a clone with the cells at the clone boundary in contact with the nearby ligand cells (as shown in Fig. 2(A)-(B)). We anticipate that the orientation of cell division with respect to the contact plane between receptor and ligand cells will play an important role. If the cell division plane is perpendicular to the receptor-ligand cell contact plane, the resulting daughter cells will each have smaller contact lengths, with both daughter cells maintaining contact with the ligand cell. On the other hand, if the division plane is parallel to the contact plane, one of the daughter cells will ‘inherit’ the full contact length while the other daughter cell will lose contact with the ligand cell. To investigate these behaviors further we quantified the time dependent behavior of cell-cell contact lengths and the cell perimeter *in silico* across multiple cell division events.

As previously described, we start at time *t* = 0*h* with a single receptor cell surrounded by ligand cells and evolve the tissue to *t* = 80*h*, whereby the receptor clone has proliferated to ~ 100+ cells. We consider two different values of the output decay rate *D* = 0 5 *h*^−1^ and *D* = 0. while keeping the synthesis rate constant. To quantify how output heterogeneity can emerge due to variations in cell properties, we focus in particular on two quantities the receptor cell perimeter, *P*_*α*_ (Fig. 3(A)-(B) and the contact length between the receptor cell *α* and the ligand cell *β* (*l*_*αβ*_), Fig. 3(C)-(D). We identified all cells located at the receptor clone interface and directly in contact with the ligand cells and recorded the values of *l*_*αβ*_ and *P*_*α*_ between *t* = 50 *h* and *t* = 80 *h*. This time interval allows us to evaluate the dynamics of *l*_*αβ*_ and *P*_*α*_ in between and across at least 2 cell division events. The gray bars denote cell division events in Figs. 3(A)-(D).

**Figure 3.**
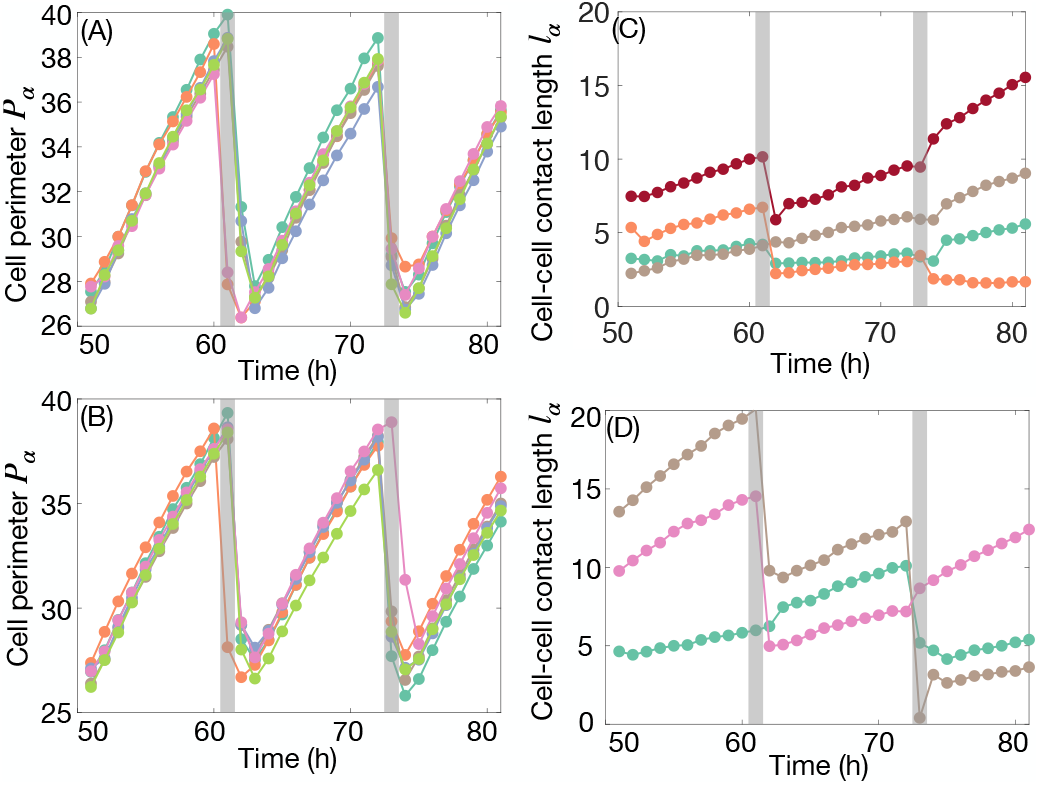
Time-dependent behavior of cell perimeter and cell-cell contact length of individual receptor cells at the periphery. (A-B) Between any two round cell divisions (marked by gray), for both (A) *D* = 0 h^−1^ and (B) *D* = 0.5 h^−1^, the perimeter (*P*_*α*_) of a border receptor cell grows linearly. Upon cell division, the perimeter of a cell drops. This sawtooth-like time-dependent behavior of the cell perimeter is highly robust and independent of intracellular output dynamics, such output synthesis *S* and output degradation *D*. Each colored dot-line represents an individual cell. (C-D) The cell-cell contact length (*l*_*α*_) of the individual receptor cells at the clone periphery, for both (C) *D* = 0 h^−1^ and (D) *D* = 0.5 h^−1^, shows highly heterogeneous time-dependent behaviors. Each colored dot-line represents an individual cell. In general, Σ_*β*_*l*_*αβ*_ = *l*_*α*_ grows linearly between any two rounds of cell division. However, the time-dependent behavior of *l*_*α*_ the instant after cell division can be broadly categorized into two different categories: (i) continues to grow linearly, or (ii) abruptly decreases. These behaviors are independent of intracellular output dynamics. Cell division events are marked by thick gray lines.

Our results show that *P*_*α*_ increases monotonically with time between any two cell division events, Fig. 3(A)-(B) (at *D* = 0 *h*^−1^ and *D* = 0.5 *h*^−1^ respectively). This is followed by a decrease in the cell perimeter during cell division, after which it increases prior to the next cell division event (see between the gray bars in Fig. 3(A)-(B). The observed time-dependent behaviors for *l* and *P* can be understood from our model. Between two successive cell division events, cell area growth is implemented by increasing the preferred area *A*_0_ at a constant rate *g*_0_ according to the equation,

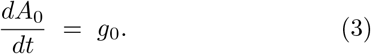

Upon cell division, a cell is replaced by two new daughter cells where we assume that cell divisions perfectly conserve the cell area. The cell area, in two dimensions of the dividing cell is equally distributed among the two resulting daughter cells,

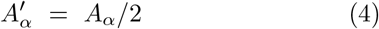

where, *A*_*α*_ is the area of cell *α* right before its division and 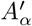 is the area of cell *α* right after division. Note that after cell division, only one of the two daughter cells carries the same index *α* as the dividing cell. A simple calculation shows that although the area is reduced by a factor of 2 immediately after cell division, the perimeter of the daughter cell is reduced by a factor of 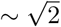,

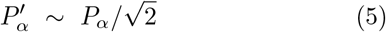

where, 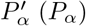 are the cell perimeter right before (after) cell division respectively. Our simulation results confirm this behavior, where the perimeter of dividing cells drops from *P* ~ 38 *µ*m to ~ 28 *µ*m, Fig. 3(A)-(B). We note that unless the cell division events are asymmetric, i.e. with daughter cells taking on different areas after division, we expect juxtacrine signals that depend on cell perimeter to lead to highly consistent time-dependent behaviors across cell division cycles.

The time-dependent dynamics of *l*_*α*_, on the other hand, is more complex. We focus on the behavior of *l*_*α*_ in two different time regimes: (i) in between, and (ii) across cell division events. In between any two cell division events, *l*_*α*_ generally increases with time (see Fig. 3(C)(D)) with the slope being highly variable from cell to cell. In Fig. 3(D), the slope for *l*_*α*_ between *t* ~ 50 − 60 h is markedly different for the three cells that are shown.

Similar to the behavior of the cell-cell contact length in between cell division events, we observe highly heterogeneous behaviors across cell division events. For example, there are instances *l*_*αβ*_ increases across cell division events (see for example gold dots in Fig. 3(C) between *t* = 50 − 70h as well as turquoise dots in Fig. 3(D) between *t* = 50 − 70h). On the other hand, there is a marked decrease in the contact length after cell division (see maroon dots in Fig. 3(C) as well as pink dots in Fig. 3(D) after the first gray bar).

Our simulation results show that when a cell divides, the magnitude of the cell-cell contact length *l*_*αβ*_ between receptor cells and ligand cells can increase or decrease immediately after cell division. We anticipate that the outcome, whether *l*_*αβ*_ increases or decreases immediately after cell division, will be determined by the cell division axis with respect to the interface between receptor and ligand cells. If the cell division axis is nearly perpendicular to the cell-cell contact length, we anticipate *l*_*αβ*_ to decrease after cell division. On the other hand, if the cell division axis is parallel to the cell-cell contact length, we expect *l*_*αβ*_ to increase.

## V. TIME-DEPENDENT BEHAVIOR OF THE RATIO OF CELL-CELL CONTACT LENGTH TO PERIMETER DETERMINES THE OUTPUT

The perimeter of receptor cells at the periphery of the clone exhibits a highly conserved sawtooth-like temporal behavior characterized by linear growth between two successive cell division events and a sudden drop immediately after cell division. On the other hand, the cell-cell contact length of these cells display distinctly different temporal behavior. Our results show that immediately after cell division, *l*_*αβ*_ continues to increase or could drop to a lower value. In our model, output synthesis in the receptor cells depend not only on the synthesis rate *S*, but also on the ratio 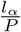, Eq. 1. Because both *P*_*α*_ and *l*_*α*_ grow linearly between two successive cell division events, we expect 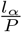 to have a constant value between two successive cell divisions, as confirmed by our simulations, Fig. 4(A)-(B).

**Figure 4.**
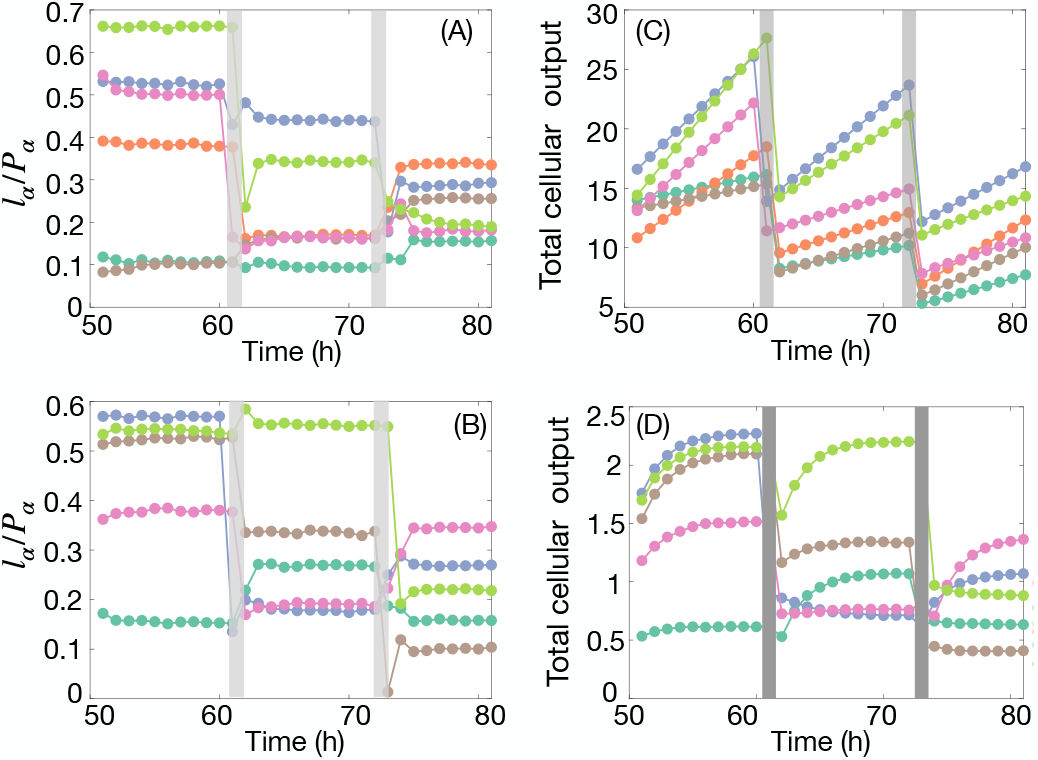
Impact of the ratio of cell-cell contact length to perimeter on the output. The ratio 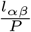 is approximately constant between two rounds of cell divisions. The total intracellular output at any time depends on the value of the ratio 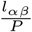. The ratio show a time-dependent behavior that is independent of intracellular output dynamics. (A) In the absence output degradation (*D* = 0 *h*^−1^) total output in individual receptor cell increases linearly with time. The slope of the linear increase of output in time depends on the ratio 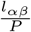. Each colored dot-line represents an individual cell. (B) However, in the presence of intracellular output degradation, total output exponentially saturates to a constant value depending on the ratio 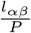. Each colored dot-line represents an individual cell. Cell division events are marked by thick gray lines.

The ratio 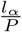 is a key parameter as the time-dependent behavior of the output depends on it. In the absence of output degradation (*D* = 0 *h*^−1^) the dynamic equation for output in a single receptor cell *α* is given by,

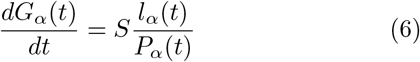

where *l*_*α*_ and *P*_*α*_, both functions of time *t*, are the total cell-cell contact length and cell perimeter, respectively. Eq. 6 shows that the dynamics of the output *G*_*α*_ depends on the time-dependent behavior of both *l*_*α*_ and *P*_*α*_. Because both *l*_*α*_ and *P*_*α*_ display a linear dependence on time between two successive cell divisions, albeit at different rates, we anticipate specific consequences on output dynamics. This suggests two things: (i) between two successive cell divisions, the quantity 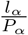 is a constant, which is given by the ratio of the rates of growth of *l* and *P*, and the output *G* has a linear dependence on time and whose rate is determined by the constant values *S* and 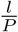. It is important to note that the parameter 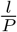 shows two distinct types of behavior immediately after cell division:

(i) step-up where the ratio increases and (ii) step-down leading to a lower value of the ratio. A step-up behavior of 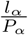 across a cell division implies a higher rate of output in the cell whereas a step-down behavior implies a lower rate of output.

We now turn to the equation describing the single cell dynamics of *G* obtained from Eq. 1 as follows:

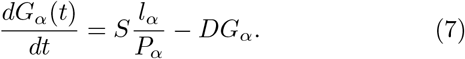

The solution to Eq. 7 is given by,

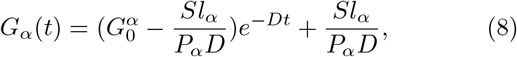

where, 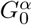 is the initial output in the receptor cell immediately after cell division. Initially, right after a cell divides, the output *G*_*α*_ grows linearly at the rate 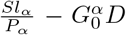. At later times, the effect of intracellular output degradation *D* is no longer negligible. This results in a gradual saturation of intracellular output *G* to a maximum value given by *G*_*max*_ = *Sl*_*α*_*/P*_*α*_*D*, which depends on the ratio of the rates of growth of contact length *l*_*α*_ and cell perimeter *P*_*α*_. Our simulation results confirm this behavior of *G*_*α*_, Fig. 4(C)-(D).

## VI. TIME-DEPENDENT BEHAVIOR OF RECEPTOR CELL-CELL CONTACT LENGTH AT THE CLONAL INTERFACE DEPENDS ON CELL DIVISION ORIENTATION

As we have seen, the output dynamics depends on the ratio 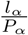 which exhibits either a step-up or a step-down behavior in time after cell division while it is approximately constant in between two cell division events. Depending on whether *l*_*α*_ continuously increases across a cell division or drops, the value of 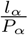 could undergo either a sudden increase (step-up) or sudden decrease (stepdown) across cell division, Fig. 4(A)-(B). We therefore wondered how cell division shapes the temporal behavior of *l*_*α*_ and *P*_*α*_ and its ratio, thus influencing the output dynamics. Our hypothesis is that the orientation of cell division with respect to the length of cell-cell contact will influence the temporal dynamics of *l*_*α*_ and *P*_*α*_ and its ratio.

To test our hypothesis, we tracked over time the receptor cells situated at the clone border through one cell division event. For our analysis, we chose the cell division event occurring at ~ 62h timepoint (see two specific cell division events highlighted in Fig. 5(A)-(B)). We chose the *t* = 50h as the initial timepoint for cell tracking whereby the receptor clone contains ~ 15-20 total receptor cells out of which ~ 10 cells are at the clone border. Cell division could lead to one of the daughter cells being pushed to the interior of the clone, leading to daughter cells that lose their position at the clone border. Therefore, in order to track the receptor cells that continue to stay at the clone border even after multiple rounds of cell divisions, we first identified all the border receptor cells at the final timepoint of the simulation, i.e., 80 hours. Each cell in the tissue has a unique index which upon cell division is randomly assigned to one of the daughter cells. Thus, we use the cell index to trace the lineage of cells in the tissue over time.

**Figure 5.**
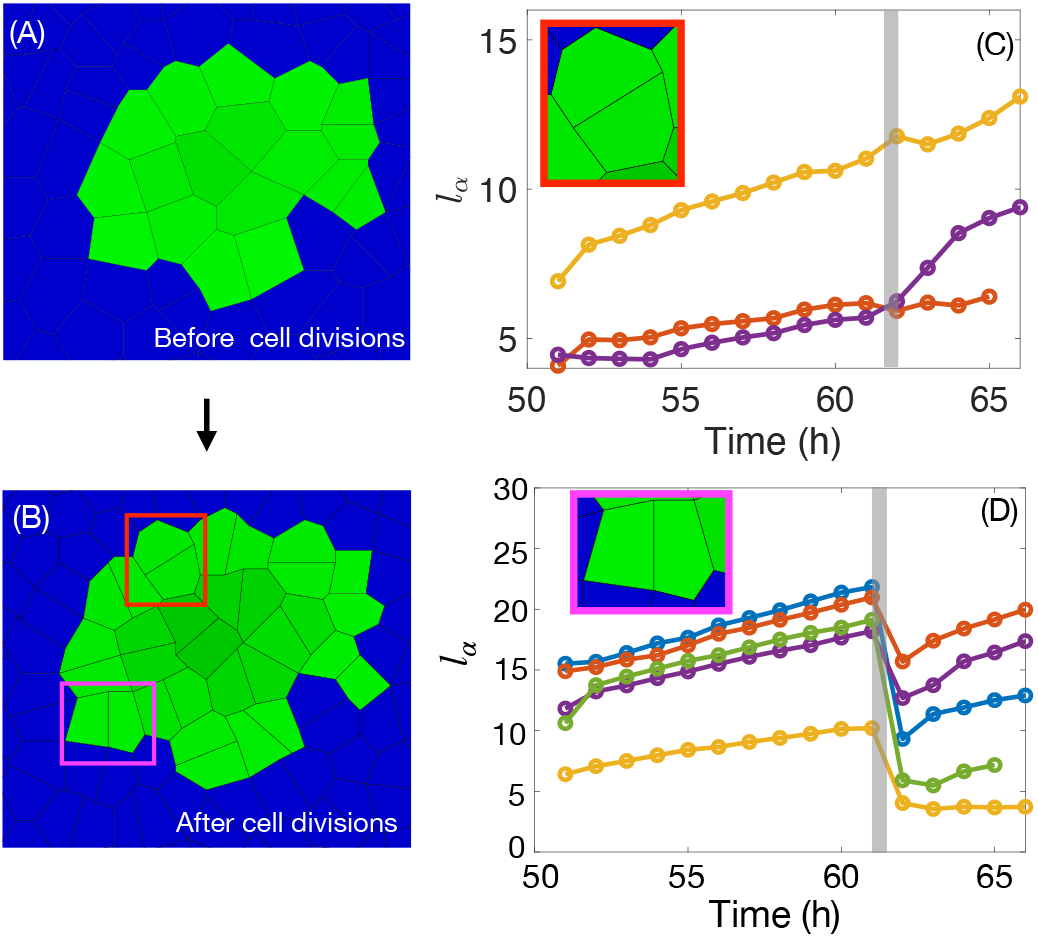
Influence of cell division orientation on the interface length of cells situated at the clonal interface. (A-B) Snapshot of the simulation showing a clone of receptor cells (green colored) surrounded by ligand cells (blue colored) (A) before cell divisions, and (B) right after cell divisions, following few rounds of cell division. Simulation results show that border receptor cells divide either parallel (red box) or perpendicular to the cell-cell contact lengths at the border (purple). Dividing border receptor cells whose division axis is parallel to the border give rise to daughter receptor cells, both of which are directly in contact with the ligand cells. However, a dividing border receptor cell whose division axis is perpendicular to the border gives rise to only one daughter cell that remains in direct contact with the ligand cells. The other daughter cell is pushed to the interior of the receptor clone. (C-D) Few dividing border receptor cells were tracked over time across one cell division (cell division event marked by thick vertical gray line). The cell-cell contact length *l*_*α*_ of the cells that divided parallel to the border (C) increases monotonically across the cell division event. However, the cell-cell contact length *l*_*α*_ of the cells that divided perpendicular to the border (D) showed a drop in their contact length right after the cell division event.

Using the cell index list of border receptor cells at the final timepoint (*t* = 80h), we sorted cell indices that are also present at the initial timepoint of interest (*t* = 50h). This filtered list contains a subset of all receptor cells that have persisted as border receptor cells through multiple rounds of cell divisions up until the final timepoint. Cells were grouped according to their division axis oriented parallel to the clone boundary as shown in Fig. 5(C) or perpendicular, see Fig. 5(D). Each border receptor cell is generally in contact with multiple cells. However, we are interested only in the dynamics of the interface cellcell contact length shared directly between a ligand and a receptor cell (green and blue cells). In particular, we are interested in how the orientation of cell division impacts the length of cell-cell contact. For a given cell *α*, the total interface cell-cell contact length is the sum of the lengths of each individual interface cell-cell contact length and is given by *l*_*α*_ = ∑_*β*_ *l*_*αβ*_(1 − *δ*_*T* (*α*),*T* (*β*)_), where *β* are neighboring cells in contact with cell *α*. The parameter *T* (*α*)(or *T* (*β*)) is the cell type value of cell *α* (or *β*) accounting for whether a cell is receptor or ligand cell. In this sum, the delta function allows us to account for contact with neighboring ligand cells and not the nearby receptor cells. Our results show that when cells that divide parallel to the clone boundary, the interface cell bond length *l*_*α*_ continues to grow across cell division. On the contrary, the cells that divide perpendicular to the clone boundary show a significant drop in *l*_*α*_. Therefore, the division axis determines the different behaviors observed in our simulations of *l*_*α*_ across cell division.

## VII. DISCUSSION

In this work, we focused on contact mediated cell-cell interaction which facilitates cell collective functions and is necessary for multicellular life [39, 40]. These cellcell contact mediated events are often highly dynamic in duration [40] and can exhibit complex cell-to-cell heterogeneity. Based on our discovery that the length of contact between signal sending ligand cells and signal receiving receptor cells largely determines the magnitude of synNotch signal output [16], we sought to understand how variations in the cell perimeter and cell-cell contact length impact signal output, especially due to cell division. We implemented a vertex-based computational model for cell growth and division coupled with contact length dependent output synthesis as well as degradation. We note that the tissue growth process leads to a highly heterogeneous output pattern within the receptor cell clones as a function of the output degradation rate *D*. The time-dependent output pattern in the entire receptor cell clone is quantified using our computational model. As the local condition and neighborhood of receptor cells may influence the output, we delved into how individual cell perimeter and cell-cell contact length varies with time, focusing on receptor cells that are actively in contact with surrounding ligand cells. While the cell perimeter exhibits a characteristic increase followed by a sharp decrease after cell division, we find that these perimeter dynamics are remarkably consistent across individual cells. However, the cell-cell contact length is highly heterogeneous and varies strongly from cell to cell. As we consider the ratio of cell-cell contact length to cell perimeter *l*_*αβ*_*/P*_*α*_together with the output synthesis rate *S*, this ratio exhibits strong variations as a function of time especially due to cell division events. After adding the contact length with neighboring ligand cells, the ratio *l*_*α*_*/P*_*α*_ for an individual cell *α* is a key determining factor in the time-dependent signal output, *G*_*α*_. We discover that the ratio is strongly dependent on the orientation of the receptor cell division with respect to the cell-cell contact length (*l*_*αβ*_). If the cell division plane is parallel to *l*_*αβ*_, the total contact length *l*_*α*_ of one of the daughter cells will increase after cell division. However, if the cell division plane is perpendicular to *l*_*αβ*_, *l*_*α*_ abruptly decreases after a division event. This leads to key differences in the time-dependent signal output associated with cell division.

Our work is focused on the pattern of juxtacrine signal output in receptor clones that are surrounded by ligand cells. This is relevant during the scenario where multiple receptor cells clustered together are surrounded by ligand cells as is observed during the development of *Drosophila* wing disc in our earlier work [16]. Other types of cell arrangement [41, 42] like the checkerboard pattern or even random arrangement of receptor and ligand cells would mean that there may not be a strong effect of cell division on the output dynamics.

### Future work

Cells are known to undergo dramatic changes in shape particularly during cell division [43–45]. How the ratio of cell-cell contact length to perimeter affects signal output levels and dynamics is not well understood. Our assumption is that the output synthesis rate is modified by the ratio *l*_*α*_*/P*_*α*_. This ratio naturally accounts for the concentration of receptors along the cell membrane. While we assume that such a concentration is uniform across all the cells this need not be the case. We plan to study how heterogeneity in the receptor concentration could affect the output levels in the future.

## ACKNOWLEDGEMENTS

AMK acknowledges support from the Jess & Mildred Fisher College of Science & Mathematics at Towson University. AMK also received support from the Faculty Academic Center of Excellence at Towson (FACET) Scholars award at Towson University. J.E.D acknowledges the support of startup grants and computing services provided by Whitworth University. J. E. D also received support from the M.J. Murdock Charitable Trust.

## VIII. APPENDIX

### 1. Vertex based modeling framework for cell shape

We use the vertex model to represent a tightly packed two dimensional cell layer. Using a cell-based computational framework, Virtual Leaf [46, 47], we simulate coupled vertex and chemical dynamics that we implemented [16]. Cells in an epithelial monolayer are represented as a network of interconnected polygons (edges as straight lines connecting vertices). The configuration of each individual cell, and thus of the whole network, is characterized by the positions of the vertices, and the connections between them. Assuming cell shape relaxations occur at timescales much faster than the cell division, the stable network configuration is determined by the minima of the energy function,

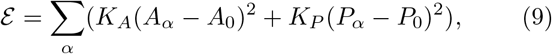

where, *A*_*α*_ and *P*_*α*_ are the area and perimeter of cell *α*, respectively. *A*_0_ and *P*_0_ are the preferred area and perimeter of cell *α*. The summation is over all cells *α* = 1 · · ·*N* (*t*), where *N* (*t*) is the total number of cells at time *t*. The first term describes the cell area elasticity with an elastic coefficient *K*_*A*_ which results from cell volume incompressibility and the cell monolayer’s resistance to height fluctuations. The second term describes the stiffness of the cell perimeter *P*_*α*_ with an elastic constant, *K*_*P*_, and it results from two processes: contractility of the actomyosin subcellular cortex and effective cell membrane tension due to cell-cell adhesion and cortical line tension. The energy function described by Eq. (9) is widely used to study tissue mechanics during morphogenesis [48, 49].

A detailed description of the VirtualLeaf vertex modeling framework, which we have adapted here, is given in Ref.[46]. The shape of a cell, and thus the whole tissue, is determined by the force balanced state, i.e., the state in which the forces, described in the energy function ℰ given by Eq.(9), acting on each node have sufficiently balanced out. The simulation of this vertex model employs Monte Carlo based ‘Metropolis dynamics’ method to alter the positions of the vertices until the energy function, given by Eq.(9), is minimized such that all interfacial tensions are balanced by cellular pressures [46]. We assume the mechanical properties of all the cells of both the cell types in the tissue, i.e., ligand and receptor-expressing synNotch cells, to be homogeneous and constant. The values of the model parameters are listed in the Table (I).

### 2. Dynamics of the output pattern in receptor cells

To characterize the spatiotemporal pattern of the output in addition to the dynamics of cell shape and cell division, we incorporate a dynamic equation for the change in the number of output molecules in a receptor cell. The output signal intensity at any time *t* in our model can be assumed to be proportional to the total number of GFP molecules in the cell at a given time. Our modeling framework thus gives rise to a coupling between three timescales at the scale of a single cell, namely, the intracellular output synthesis rate, the rate of intracellular output degradation and the rate of cell division.

In our model, we consider two types of cells, namely, synthetic ligand expressing cells and synthetic Notch receptor expressing cells (which we refer to as receptor cells). The ligand expressing cells are assigned a label or cell type of 0 and the receptor cells are assigned a label or cell type of 1. This binary-type quantity *T* is a function of cell *α*, where *α* is the cell index, *i* ∈ 1 … *N*. *T* takes the value *T* (*α*) = 0 if the cell *α* is a ligand expressing cell and *T* (*α*) = 1 for a receptor cell, Fig. 1(A). The total number of cells at any given time point is *N* (*t*) = *N*_(*T* =0)_ + *N*_(*T* =1)_, where *N*_(*T* =0(1))_ is the total number of cells of type *T* = 0(1). We assume that the entire monolayer is fully characterized as comprising of type 0 and type 1 cells. The cell identity, *T* (*α*) = 0 or 1, assigned to a cell is conserved across cell division events. Therefore, when a cell *α* of type *T* (*α*) = 0, i.e., a ligand expressing cell, divides, it produces two daughter cells, both of which are of cell type 0. Similarly, when a cell of cell type *T* (*α*) = 1, i.e., a receptor cell, divides, it produces two daughter cells, both of which are cells of cell type 1. The cell type *T* is associated with the kronecker delta function *δ*_*T* (*α*),*T* (*β*)_, which assumes the value 0 if the cell *α* and its adjacent cell *β* are of different types, i.e., (*T* (*α*) ≠ *T* (*β*)), and, the value 1 if they are of the same type, i.e., (*T* (*α*) = *T* (*β*)). With this condition, output is restricted to only those cells that share a cell-cell junction with synthetic ligand expressing cells. We propose that the output in a receptor cell is proportional to the length of the cell-cell contact length, *l*_*αβ*_ (*µ*m) shared with its adjacent synthetic ligand expressing cell, and normalized to the overall perimeter *P*_*α*_ (*µ*m) of the receptor cell *α* at time *t*.

### 3. Rules governing cell growth and division

Each cell in the *in silico* tissue is fixed to the same base area, *A*_*b*_. Our initial condition is such that all the cells are set to *A*_*α*_ = *A*_*b*_ = 50, where *α* is the cell index. At *t* = 0, the preferred area *A*_0_ of all the cells is equal to the base area. The dimensionless value of 50 for the cell area in the model corresponds to an experimental value of 10*µ*m^2^. To implement cell area growth over time, as described in [46], the cell preferred area is increased in the next time step *t* + *δt* as,

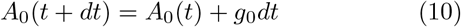

where, *g*_0_ is the value by which the cells preferred area is incremented in the next time step. The cell area at each time step is relaxed quasi-statically to the preferred area, which results in an irreversible increase in the cell area over time. Over time, when the area of a cell *A*_*α*_ is double the base area *A*_*b*_ then the cell divides over its short axis, Fig. 1(D-iv). Right after division, the preferred area of the divided cell is reset to *A*_0_ = *A*_*b*_ = 50. As a result the area of each daughter cell, the instant after cell division, is relaxed to *A*_0_ = *A*_*b*_.

Since the cell divides when its area *A*_*α*_ reaches a value which is twice the base area of the cell *A*_*b*_, which occurs in the time duration *τ*,

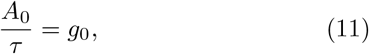

in our simulations we chose a timescale such that 1 relaxation step that minimizes the energy function *E*, given by Eq. (9), for the preferred *A*_0_(*t*) at the time *t*, corresponds to 1 h. We set *A*(*t*) = *A*_0_, where *A*_0_ = 10*µm*^2^ is the typical base area of a cell in the wing disc epithelium. The typical time *τ* over which a wing disc epithelium cell divides is equal to 10 h. Hence, *g*_0_ = 1*µm*^2^*/h* is the typical cell area growth rate. By dividing and multiplying both sides of Eq.(11) by a unit area *A*\* = 0.2*µm*^2^ and a unit time *t*\* = 1*h*, respectively, we obtain,

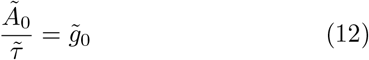

We chose the cell base area as *Ã*_0_ = 50, which corresponds to the experimental value of 10*µm*^2^, and cell division time 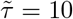, which is equivalent to 10h, implying that 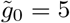.

